# Transcriptionally active chromatin loops contain both ‘active’ and ‘inactive’ histone modifications that exhibit exclusivity at the level of nucleosome clusters

**DOI:** 10.1101/2023.09.03.555774

**Authors:** Stefan A. Koestler, Madeleine L. Ball, Leila Muresan, Vineet Dinakaran, Robert White

## Abstract

Chromatin state is thought to impart regulatory function to the underlying DNA sequence. This can be established through histone modifications, and chromatin organisation, but exactly how these factors relate to one another to regulate gene expression is unclear. In this study, we have used super-resolution microscopy to image the Y loops of *Drosophila melanogaster* primary spermatocytes, which are enormous transcriptionally active chromatin fibres, each representing single transcription units that are individually resolvable in the nuclear interior. We previously found that the Y loops consist of regular clusters of nucleosomes, with an estimated median of 54 nucleosomes per cluster with wide variation. In this study, we report that the histone modifications H3K4me3, H3K27me3, and H3K36me3 are also clustered along the Y loops, with H3K4me3 more associated with diffuse chromatin compared to H3K27me3. These histone modifications form domains that can be stretches of Y loop chromatin micrometres long, or can be in short alternating domains. The different histone modifications are associated with different sizes of chromatin clusters and unique morphologies. Strikingly, a single chromatin cluster almost always only contains only one type of the histone modifications that were labelled, suggesting exclusivity, and therefore regulation at the level of individual chromatin clusters. The active mark H3K36me3 is more associated with actively elongating RNA polymerase II than H3K27me3, with polymerase often appearing on what are assumed to be looping regions on the periphery of chromatin clusters. These results provide a foundation for understanding the relationship between chromatin state, chromatin organisation, and transcription regulation – with potential implications for pause-release dynamics, splicing complex organisation and chromatin dynamics during polymerase progression along a gene.

## Introduction

Cells must undergo tightly regulated patterns of differentiation, as well as maintain the ability to dynamically respond to external stimuli. These functions are underpinned by dense genetic networks, with specific regulation of gene expression crucial to both cellular identities and responses. This precise genetic regulation is thought to be achieved through a diverse range of transcription factors, epigenetic modifications, and chromatin organisation that control levels of transcription (1). However, it is still mostly unclear exactly how these different elements work together. The genome is organised at different levels within the nucleus: into individual chromosomes, that occupy spatially distinct chromosome territories (2–4), different active and inactive domains visualised as euchromatin and heterochromatin (5), or as A and B compartments (6,7), and further into topologically associated domains (TADs) (8,9), and loops (10,11). At the finest level of organisation, the DNA of the genome is packaged into chromatin, which is formed through the complexing of DNA with histones to form nucleosomes. This compacts the helical DNA fibre and allows for the formation of higher-order domains. The chromatin landscape can be epigenetically modified and acts as a “landing pad” for many chromatin-associated proteins that can regulate gene expression, particularly interacting with post-translational modifications like methylation and acetylation of the N-terminal tails of the histone proteins (12–14).

A key feature of the nucleosome is the dynamic nature of the association with DNA, with DNA being capable of sliding significant distances in relation to the core complex, as well as tightening or relaxing the association to create denser or looser fibres (15,16). This constant deformation of the nucleosomes has been referred to as ‘breathing’ motions, and contributes to the liquid-like, dynamic organisation of chromatin (17,18). The relaxed association of the DNA fibre to the core histone complex is thought to allow access for transcription factors to bind their target DNA sequence and subsequently lead to the activation of transcription, whereas the tighter association of DNA around the histone core complex is thought to block these binding interactions, therefore preventing transcription from occurring at these loci (19). During elongation RNA polymerase also has to overcome the nucleosomal barrier to progress along the template DNA, requiring dynamic replacement, displacement, and repositioning of nucleosomes (20,21). Histone post-translational modifications can recruit factors such as chromatin remodelers that can directly cause these structural changes to chromatin by dynamically repositioning, removing, or exchanging nucleosomes in an ATP-dependent manner (22). Many chromatin remodelling complexes have been identified that are essential for development. One example is the ISWI (imitation switch) family of complexes that are important for regulating high-order chromatin structure – knockouts of ISWI in *Drosophila* lead to a dramatic decondensation of chromatin (23,24). This interaction between epigenetic modification, recruitment of factors and the subsequent modification to chromatin structure thus provides a mechanism through which histone modifications can impart their regulatory effect.

Histone modifications can be broadly split into two categories; ‘active’ marks and ‘inactive’ or repressive marks, that are associated with transcriptionally active and transcriptionally repressed regions of the genome respectively. These histone modifications have been associated with different chromatin states, both inactive and active marks correlate with different chromatin structural organisations that are thought to work alongside other factors to regulate transcription. Active and inactive regions of the genome tend to cluster in the genome linearly as shown by ChIP data (25,26), but based on Hi-C and 3C-based methods, they also cluster spatially into 3D domains (6). These domains of active and inactive chromatin modifications correlate to regions of high and low transcriptional activity respectively. However, the details of mechanistic links between chromatin state and gene expression are still not fully understood. Super-resolution microscopy analysis has begun to link the structural organisation of chromatin and different chromatin states, for example by showing that chromatin modified with active marks have smaller, spatially separated clusters, and denser chromatin being associated with inactive marks (27,28). Also super-resolution analysis on mouse pachytene chromosomes revealed specific organisation of chromatin clusters associated with different histone modifications (29). Using a combination of Hi-C, FISH, and super-resolution microscopy in *Drosophila* nuclei, repressed TADs were shown to be organised into condensed physical entities, or nanodomains, while active domains consisted of more open, decondensed chromatin – with the two types of structure feature being interspersed with each other (30).

To what extent chromatin structure regulates gene expression is still a matter of debate. It has previously been shown that even highly rearranged balancer chromosomes are not often associated with disruptions of gene expression (31), and that highly condensed nucleosome domains are found even in active regions (32). Super-resolution microscopy analysis can be utilised to image chromatin organisation directly in single cells, however, this is usually hampered by the dense packing of chromatin in most nuclei, making distinguishing different chromatin fibres from one another a significant challenge. To overcome this obstacle, we have exploited the large nuclei of *Drosophila melanogaster* primary spermatocytes and their Y loops; single fibre, transcriptionally active chromatin loops that emerge from the Y chromosome as several megabase long structures. We have previously visualised and quantified the chromatin structure of the Y loops at super-resolution, showing that they are organised as chains of nucleosome clusters, with an average cluster width of approximately 50 nm (33). This aligns with current opinion based on multiple complementary techniques that consider interphase chromatin *in vivo* to be arranged as a heterogenous, discontinuous structure of irregular clusters or ‘clutches’ as opposed to a continuous 30 nm fibre (34). Here we take advantage of the Y loops as a model system to visualise and quantify the organisation of chromatin state along single transcriptionally active fibres.

We find that the chromatin of the Y loops is not homogenous with respect to chromatin state and these active loops contain a mix of domains carrying either active or inactive marks. There is considerable variety in the arrangements of these chromatin state domains along active chromatin fibres, presumably reflecting the dynamics of the various transcriptional processes. However, at the level of nucleosome clusters the examined histone marks show exclusivity which, we suggest, reveals a key relationship between the organisation of chromatin into nucleosome clusters and chromatin state.

## Results

### Histone modification distribution along Y loops

In *Drosophila* spermatocytes, as part of the spermatogenesis transcription program, three extraordinarily large genes on the Y chromosome become activated and expand into the nucleoplasm as visible Y loops. Each loop comprises a single transcription unit, several megabases in length, and these Y loops provide a useful model for the visualisation of actively transcribed chromatin (35,36). Here we are interested in the distribution of histone modifications within these transcription loops to investigate how histone modification is linked to variation in chromatin structure. For this, we first examined the distribution of one of the key modifications of active chromatin, H3K4me3 (37) by stimulated emission depletion (STED) microscopy (38) on isolated primary spermatocytes immuno-labelled with pan-histone and H3K4me3 specific antibodies (Fig. 1A; the pan-histone antibody labels the core histones and H1 and we will refer to it hereafter as simply histone labelling). To aid overall visual inspection, signal to noise was improved by denoising with N2V (see Materials and Methods), but quantifications were performed on the raw data. H3K4me3 labelling was most apparent in chains of clusters close to the nucleolus, with more sparse labelling of loops in the central nucleoplasm (Fig. 1). Some long Y loop regions lack apparent H3K4me3. Notably, the H3K4me3 labelling intensity generally does not directly follow the histone labelling density and the peaks of H3K4me3 intensity overlay rather weak histone labelling, suggesting that the H3K4 mark is associated with relatively disperse chromatin (quantification is presented below). Although the H3K4me3 mark has been specifically linked to transcription initiation (39), the Y loop labelling is clearly not tightly restricted to the sites of initiation which are thought to be close to the Y chromosome mass associated with the nucleolus (33). Instead, the chains of H3K4me3 labelled clusters emanating from the nucleolus indicate that the H3K4me3 mark extends well into the transcription unit. This and the labelling further along the body of the Y loop transcription units would be consistent with the recent finding that H3K4me3 has a distinct role in transcriptional pause-release and elongation rather than transcriptional initiation (40).

**Figure 1:**
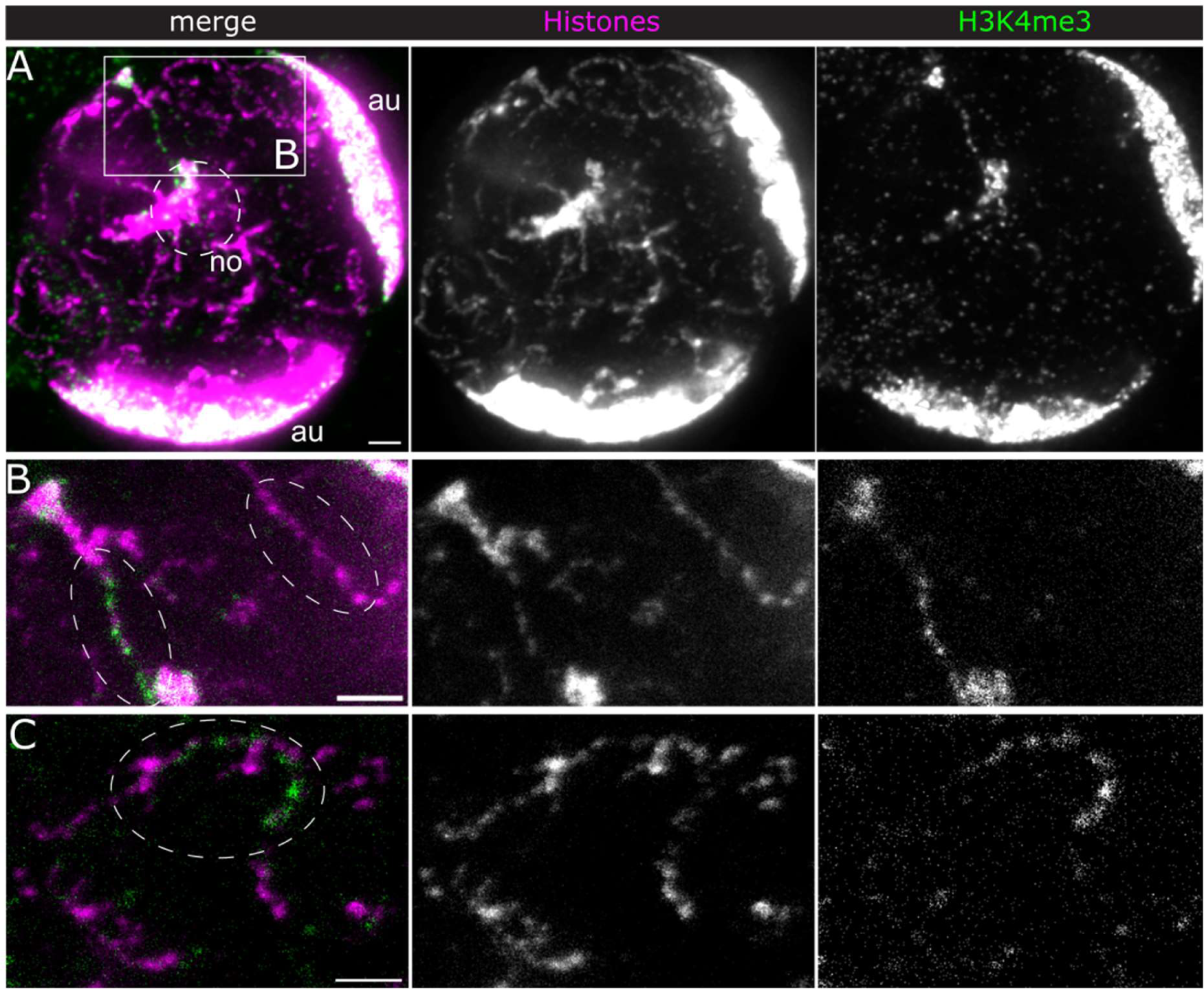
H3K4me3 distribution along Y loops. STED immunofluorescence (IF) microscopy of histones (magenta and central panel) and H3K4me3 (green and right panel). A) overview of a spermatocyte nucleus, N2V denoised and maximum projected over 5 µm (10 slices with Δz=0.5 µm). no, nucleolus with sex chromosomes; au,dense autosome masses at the nuclear periphery. B) Detail of boxed region in A) showing a single optical unfiltered (raw) section of a Y loop fibre enriched with H3K4me3 and one without H3K4me3 nearby (ellipses). C) Example of a fibre with one H3K4me3 enriched segment in another cell. Maximum projection of 4 unfiltered (raw) slices. Of note, the same STED laser was used for both excitation lasers, thus eliminating the potential for any artificial shifts between the two fluorophores due to misalignment. Scale bars, 1 µm.

We then examined the distribution of H3K36me3, a well characterised marker for active transcription along gene bodies (41). We expected H3K36me3 to be enriched along Y loops, since they function as single transcription units (35,36). However, we only found high signals at a prominent “spike” region protruding from the nucleolus into the nucleoplasm (Fig. 2A) and rather sparsely along fibres in the nucleoplasm (Fig. 2B). Fig. 2C shows an example of H3K36me3 labelling peripheral to the major histone density along the fibre, suggesting that H3K36me3 is associated with relatively decondensed chromatin. The punctate H3K36me3 signals along fibres in the nucleoplasm may indicate the dynamics of the H3K36me3 mark.

**Figure 2:**
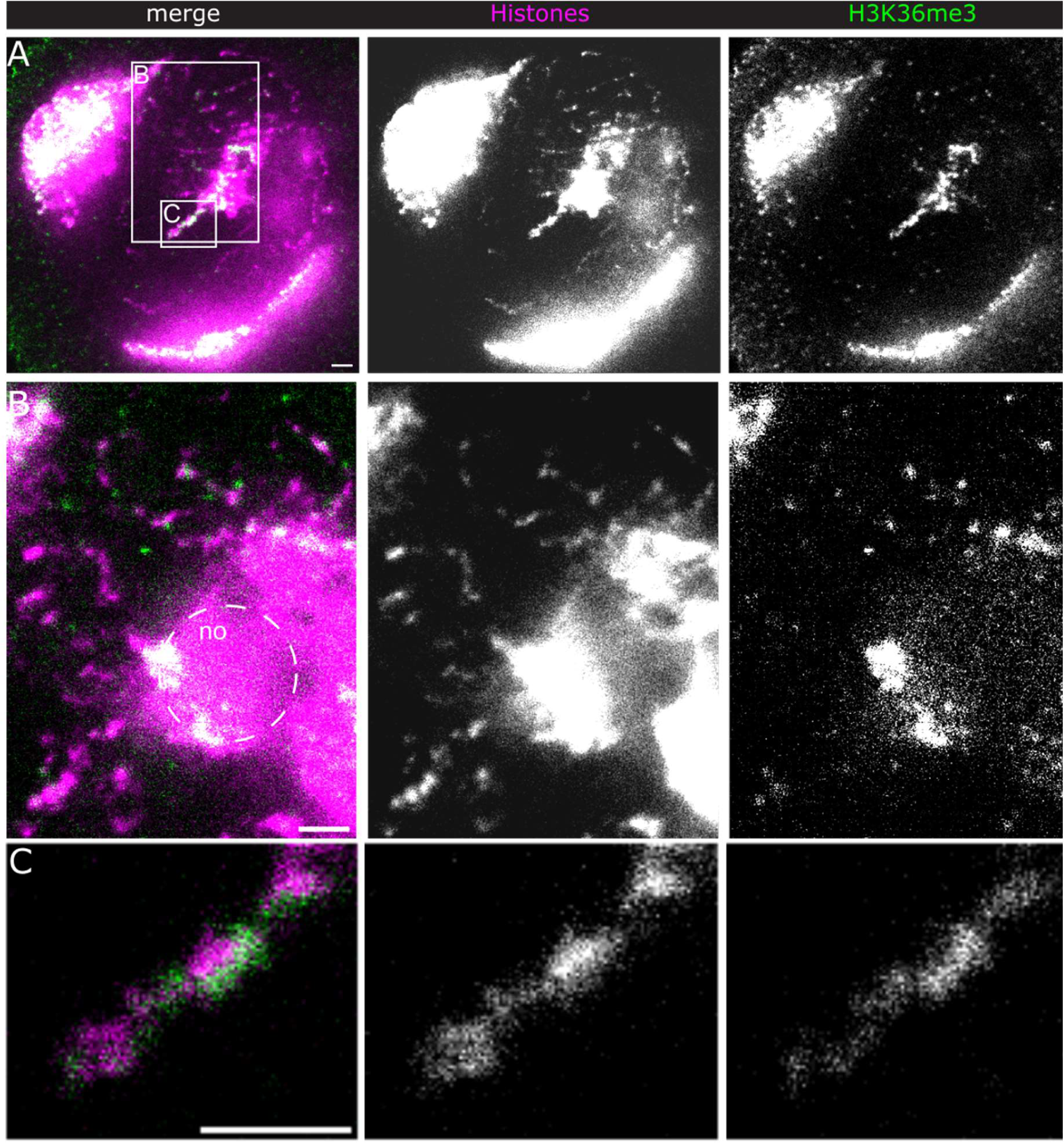
H3K36me3 distribution. STED IF microscopy of histones (magenta and middle panels) and H3K36me3 (green and right panels). A) overview of a spermatocyte nucleus (single slice). B)&C) details of boxed regions in A). B) shows fibres in the nucleoplasm and part of the nucleolus. C) “Spike” region emanating from the nucleolus. All unprocessed data. Scale bars, 1 µm.

The observation that the accumulation of active chromatin marks, especially H3K4me3, is highly restricted along Y loops prompted us to determine if we could detect a mark that is associated with inactive transcription. Hence, we turned to H3K27me3, which is a well-established marker of the repressed chromatin state (42). Surprisingly, STED microscopy revealed a prominent clustered labelling of H3K27me3 along many, but not all, Y loop regions (Fig. 3A). Comparison of the H3K27me3 and histone labelling densities (Fig. 3B) suggests that H3K27me3 is associated with both condensed and relatively decondensed chromatin.

**Figure 3:**
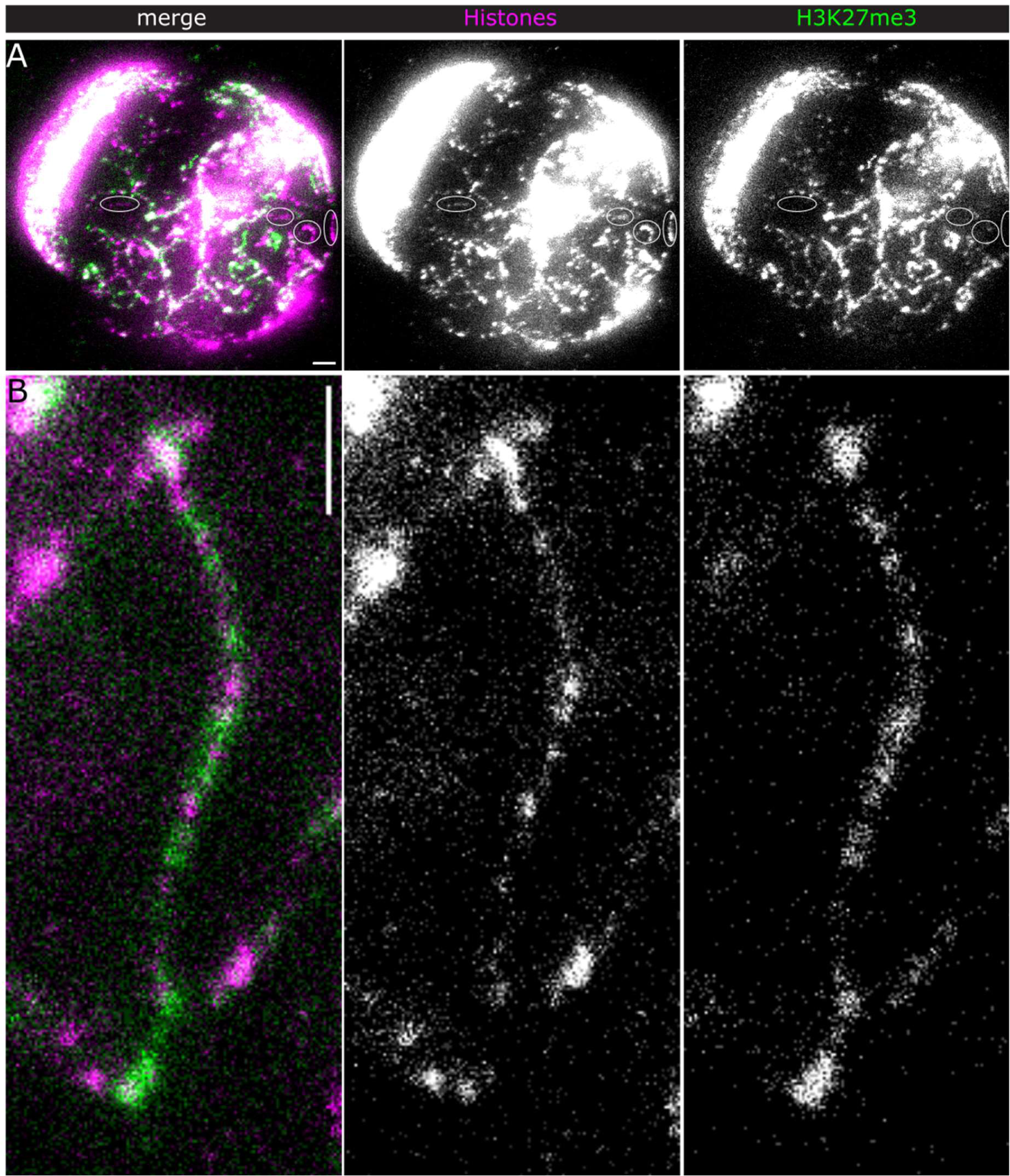
H3K27me3 distribution along Y loops. STED IF microscopy of histones (magenta and middle panels) and H3K27me3 (green and right panels). A) overview. Ellipses indicate fibres without detectable H3K27me3. Maximum intensity projection of 10 slices, Δz = 0.5 µm. B) shows a Y loop fibre with H3K27me3 clusters in decondensed chromatin regions. Maximum intensity projection of 4 slices, Δz = 0.5 µm. Scale bars, 1 µm.

### Direct comparison between H3K27me3 and H3K4me3 labelling

Each Y loop represents a single gene that is transcribed throughout its whole length and the transcriptional processes must be coordinated. Thus, the relative distributions of marks for different transcriptional activities can be expected to be informative about their mechanism and dynamics. Co-labelling of H3K4me3 and H3K27me3 and observation by laser scanning confocal (LSC) and STED microscopy showed that labelling of these two marks is mutually exclusive (Fig. 4). This occurs however in various arrangements, from alternating clusters to long exclusive fibre sections of either mark (Fig. 4B). This indicates that the transcriptional dynamics along the Y loops are highly variable including a high turn-over of histone modifications and chromatin rearrangements. In some instances the opposing histone marks occur along the same section of the fibre, not overlapping however, but with H3K27me3 more central and H3K4me3 more distal to the main fibre axis (Fig. 4E). This suggests a mechanism for transcription off the main fibre, from which low density chromatin, might be looping out and potentially spread to form the more extended transcriptionally active chromatin.

**Figure 4:**
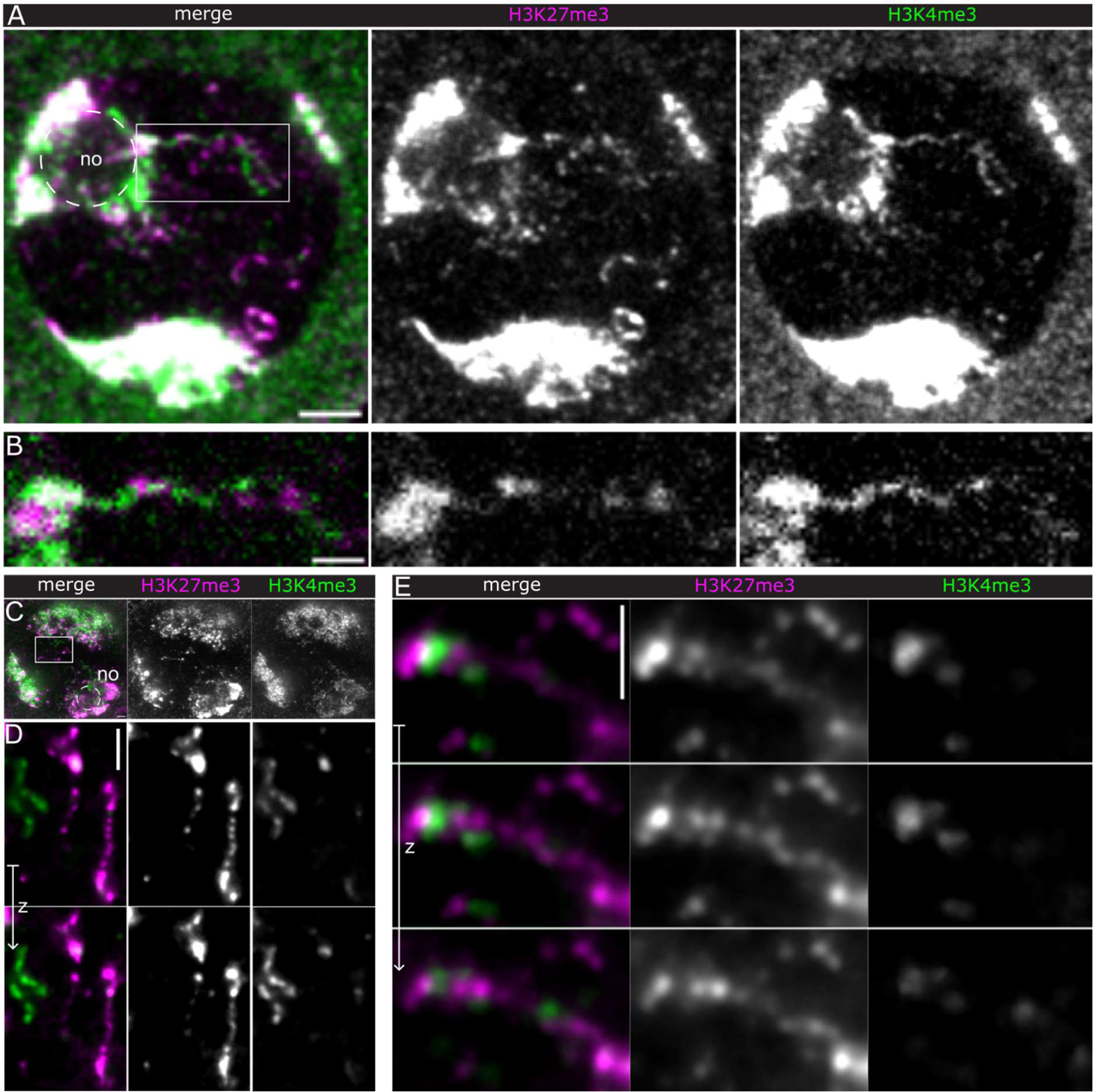
H3K4me3 and H3K27me3 distributions. A)&B) LSCM of H3K4me3 and H3K27me3 labelled spermatocytes. The ImageJ smooth filter was applied. A) overview, maximum intensity projection of 4 slices (Δ z = 300 nm). Bar, 3 μm. B) Detail of boxed region in A) showing a fibre emanating from the nucleolus and carrying both histone marks. Single optical section. C)-E): N2V denoised STED images. C) overview. D)&E) details, 3 optical sections (Δ z = 450 nm). D) shows fibres in the same nucleus almost exclusively stained either for H3K4me3 or H3K27me3. Images are rotated by 90° relative to C). E) Detail from another cell showing H3K27me3 clusters along a fibre with locally associated H3K4me3. Scale bars, 1 µm.

### Quantification of histone modification distributions

To investigate the link between chromatin architecture and active and inactive chromatin marks along Y loops we first measured the distances between histone clusters on long (µm range) fibre sections enriched either with H3K4me3 or H3K27me3 (Fig. 5 A-C). Since higher chromatin densities are often thought to impede efficient transcription and also have been shown to stimulate PRC2 activity (43) and lower transcriptional activity corresponds with high H3K27me3 levels, we expected fewer clusters and more extended chromatin in regions displaying the active H3K4me3 mark. Strikingly, there was no difference in the distance between histone clusters, with a mean of 276+-164 nm for H3K4me3 decorated fibres versus 271 +-157 nm for fibres enriched with H3K27me3 when applying a low tolerance factor for peak detection, and 123+-52 nm vs 126+-58 nm with a low stringency.

**Figure 5:**
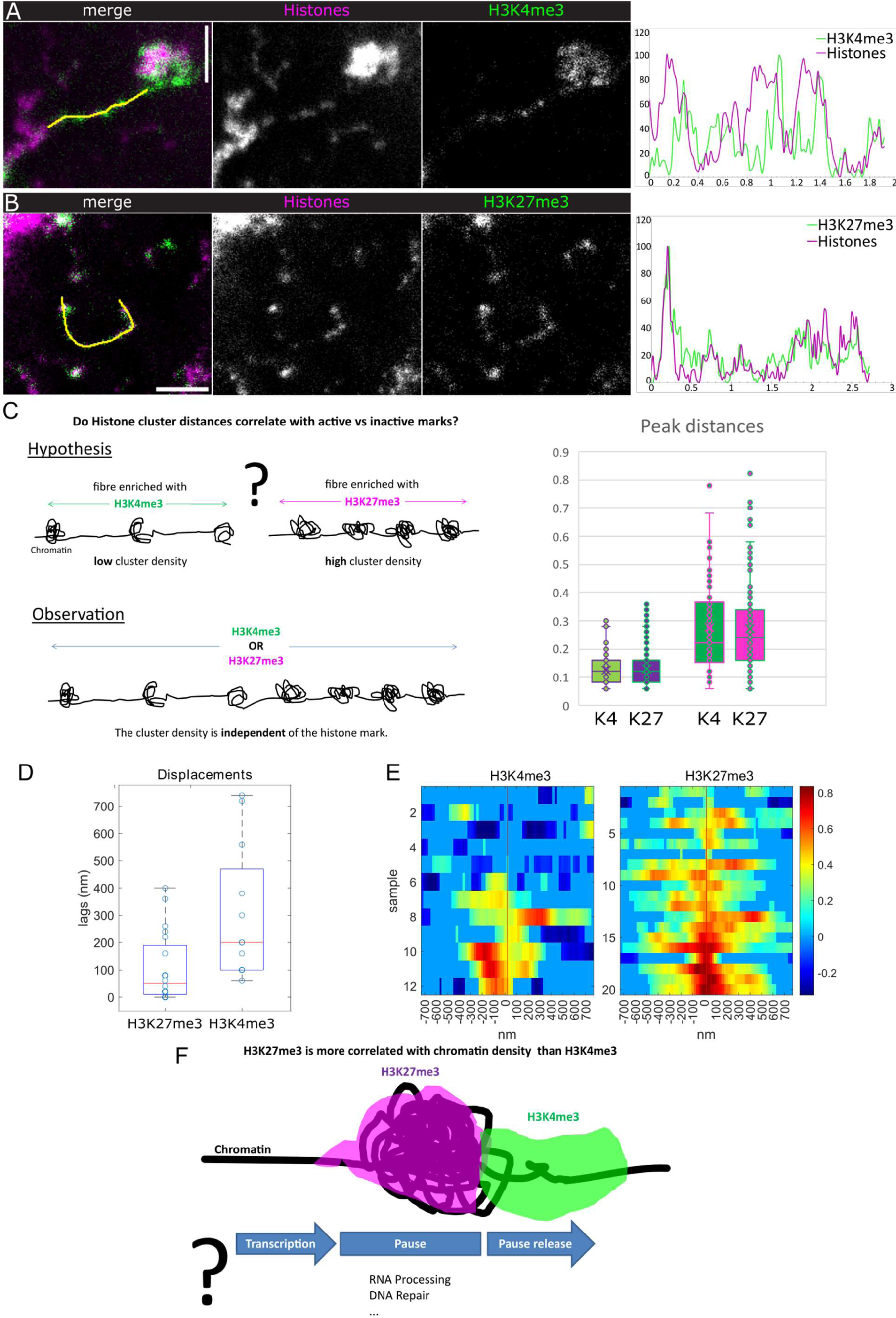
Quantitative analysis of the relationship between histone modifications and chromatin densities. Corresponds to data as shown in Figs. 1&3. A)&B) right panels show example intensity profiles along the indicated line in the left (merge) panels. X-axis of the plots gives the distance in µm, normalized intensities are along the y-axis. A) H3K4me3 vs histones, example from Fig. 1. B) H3K27me3 vs histones. C) The scheme indicates the expected distribution of chromatin clusters in H3K4me3 vs H3K27me3 enriched Y loop regions, and the distributions as observed by measuring the distances between histone intensity peaks. Quantifications are given in the box plots; left, for a tolerance factor for peak detection of 0.17, right, for a tolerance factor of 1.0 to avoid a bias caused by false positive or negative detections. Number of measured peak distances are 169, 266, 54 and 102. Total measured Y loop lengths are 23.42 µm (H3K4me3) and 37.76 µm (H3K27me3). D)&E) The cross correlation of fluorescence signal intensities along selected loops was computed for H3K27me3 vs histones and H3K4me3 vs histones respectively. D) shows the overall displacement distributions in box plots. E) Heat maps indicate the cross correlation values between the lagged signals of respective cluster intensities per nucleus. The x-axis shows the lag distances, the samples are stacked along y. F) The scheme indicates the shift between histones and the respective histone marks, as well as their mutually exclusive organization (see Fig. 4). The question mark indicates our interpretation of the results that H3K27me3 enrichment at chromatin clusters is correlated with transcriptional pause, which would allow e.g. processing of the transcribed RNA or DNA repair to occur. H3K4me3 enrichment at the periphery of the clusters could indicate pause release into productive elongation.

This shows that the basic clustered chromatin organization is independent of the studied histone marks, and even regions carrying active marks contain chromatin clusters suggesting that this organization is permissive for transcription.

To obtain a more direct measure of the relationship of H3K4me3 and H3K27me3 with chromatin densities we calculated their respective cross correlations (Fig. 5A, B, D, E). We find a stronger direct correlation of chromatin with H3K27me3, while H3K4me3 is usually found displaced from the chromatin enrichments (median absolute value of shifts (displacement): 50 nm for H3K27me3 and 200 nm for H3K4me3). The t-test for the absolute shift gives a p-value of 0.0049.

Thus, while these two histone marks do not appear to affect chromatin cluster frequencies along Y loops, their differential enrichment with respect to chromatin densities suggest transcription dynamics with inactive transcription or pausing at clusters and transcription progression at their periphery (44). Pausing could allow e.g. DNA repair or RNA processing, which might form a particular globular chromatin arrangement (Fig. 5F).

### Different chromatin states have different structural organisation

To extend this analysis and to assess the morphology of chromatin in different activity states we took advantage of the higher resolution of STORM and imaged Y loops single labelled for H3K36me3, H3K4me3 or H3K27me3 (Fig. 6).

**Figure 6:**
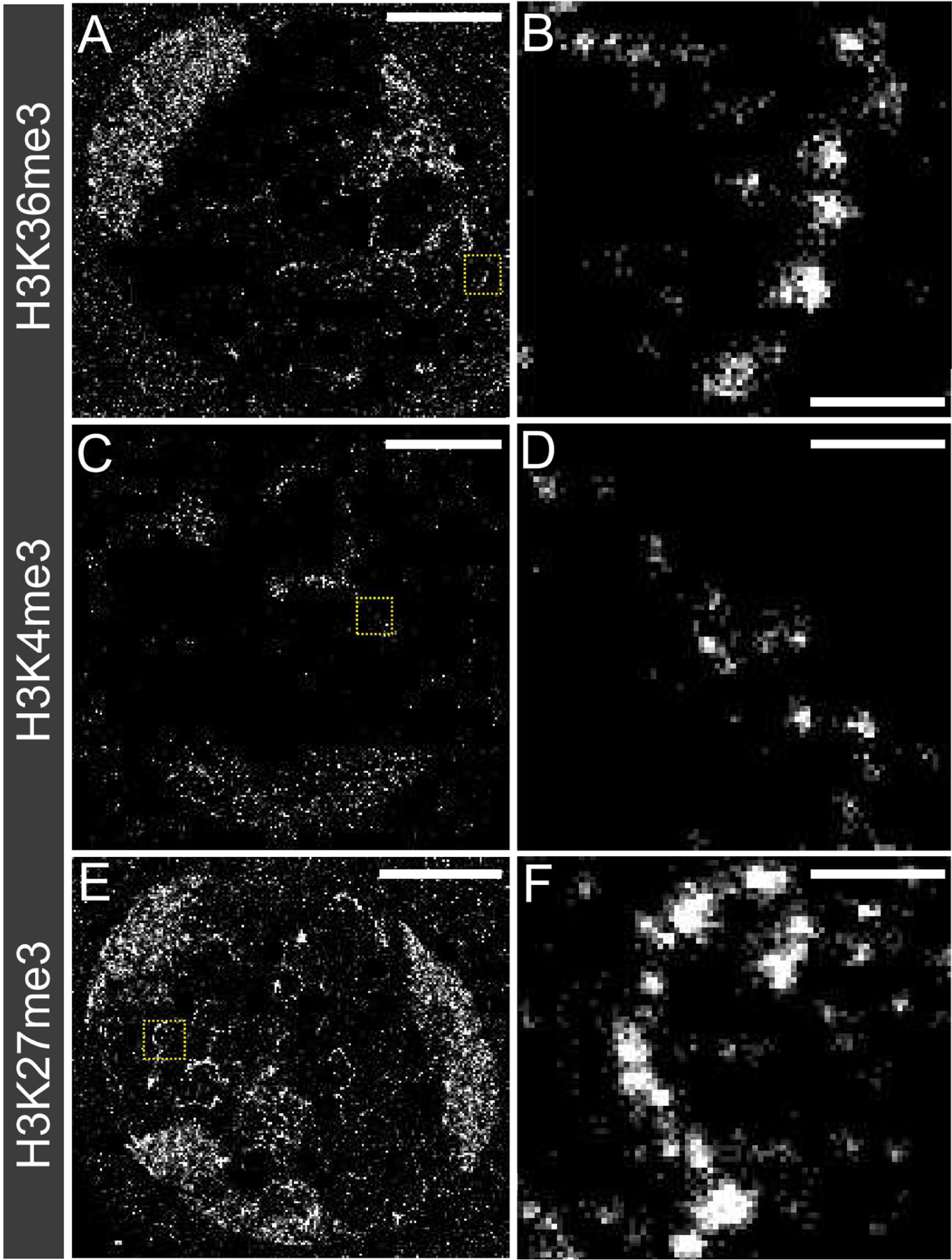
Single-labelled super-resolution STORM images of primary spermatocyte nuclei. A&B) labelled for H3K36me3, C&D) labelled for H3K4me3 and E&F) labelled for H3K27me3. The scale bars for left hand panels are 5 µm. Zoomed in images of yellow box areas are shown on the right. The scale bars in the zoomed in images are 500 nm.

The chromatin clusters labelled for H3K36me3 show heterogenous morphology. The clusters appear to be larger than global histone labelled Y loop clusters (33), with more “fuzzy” edges, suggesting the presence of looping DNA emanating from the edges of the clusters. Interestingly, there appear to be clear runs of Y loop chromatin which have continuous enrichment for H3K36me3, as the clusters appear approximately 100 nm apart from one another, the same spacing as the general histone labelled Y loop chromatin. We previously estimated that the average chromatin cluster along the Y loops could contain a median of 54 nucleosomes. In terms of sequence this means that a cluster could be formed of approximately 8.2 kb, and therefore continuous clusters along a fibre modified for the same histone modification implies that very large domains of H3K36me3 modified chromatin are present along the Y loops.

The chromatin clusters of H3K4me3 visually appear much smaller when compared to the Y loops labelled with the histone antibody, and the Y loops labelled for H3K36me3. The H3K4me3 clusters also appear to be more disconnected from each other (Fig. 6D) suggesting that there may be intervening clusters of nucleosomes that do not have the H3K4me3 modification, or that H3K4me3 is enriched in the regions in between the chromatin clusters of the Y loops. This suggests that there are not clear continuous domains of H3K4me3 along the Y loops as was the case for H3K36me3.

H3K27me3 appears to represent mostly large clusters along the Y loops, visually appearing similar to H3K36me3. However, the H3K27me3 labelled chromatin clusters appear to have sharper boundaries suggesting they lack the same extent of decondensed “looping” structure that surrounds the H3K36me3 clusters (Fig. 6D). There are also continuous runs of clusters of chromatin along fibres enriched for H3K27me3, implying the existence of large domains (Fig. 6F). We note that H3K27me3 is not exclusively found labelling larger clusters, but rather can be seen labelling a wide variety of different cluster sizes along the Y loops.

The chromatin clusters labelled by H3K4me3, H3K36me3 and H3K27me3 visualised by STORM therefore showed unique morphologies.

### Quantification of structural organisation

In order to better characterise the structural differences of chromatin modified for active and silencing histone modifications, clustering analysis was performed as described previously (33). This analysis measures the median width (full width half maximum; FWHM) of the clusters identified, enabling the comparison of clusters of chromatin in different states (Fig. 7). Due to the variable numbers of clusters used in each analysis, the frequencies were normalised, and presented as a ‘probability’ value instead of raw frequency counts on the histogram.

**Figure 7:**
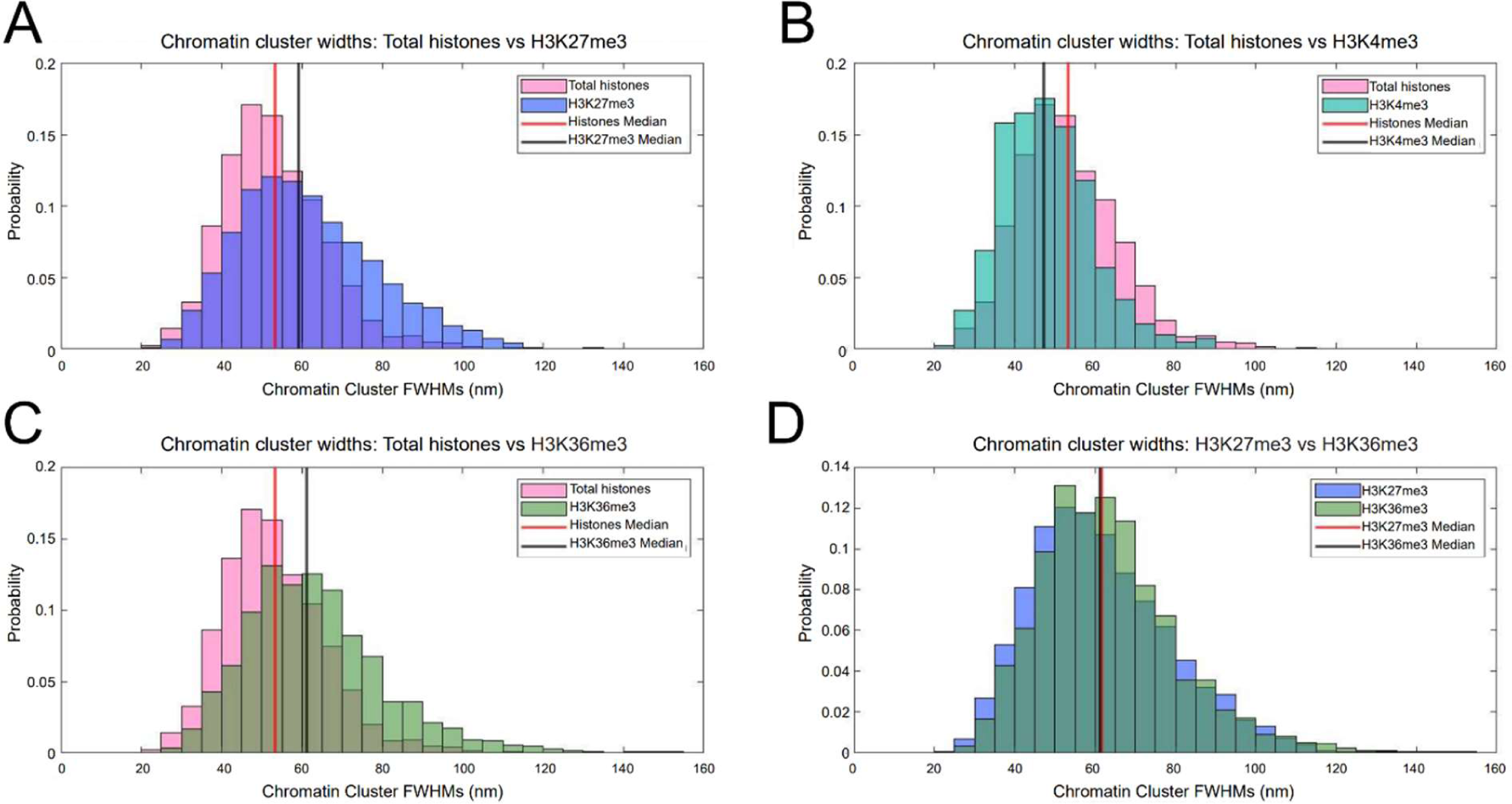
Histograms showing the cluster widths (FWHM) of Y loop clusters labelled with different histone modifications. Total histone labelled compared to A) H3K27me3, B) H3K4me3, C) H3K36me3, and D) H3K37me3 compared to H3K36me3. For A), B), and C), the red line indicates the median cluster width of total histones, and the black line indicates the median cluster widths of the histone modification. For D), the red line indicates the median cluster widths of H3K27me3 and the black line indicates H3K36me3.

The histone labelled chromatin cluster widths are as described previously (33), with a median of 52 nm, and an interquartile range (IQR) between 44 and 61 nm. H3K27me3 labelled chromatin had a median width of 59 nm, and an IQR between 49 and 72 nm. Compared to the general histone label, the H3K27me3 modified chromatin showed a bias towards larger clusters, with approximately a 14% increase in median width (P = <0.05). It is important to note that all the modified chromatin cluster sizes were almost exactly within the range of the general histone data, and so the cluster sizes are not larger or smaller than what can be seen in the general histone data. However, the skew of the graphs was different, indicating a bias towards finding different chromatin cluster sizes depending on histone modification. The H3K36me3 labelled chromatin clusters had a median width of 61 nm, and an IQR between 51 nm and 72 nm. This represents an 18% increase in median width compared to global Y loop histone labelling (P = <0.05). The H3K4me3 appeared visually much smaller by eye, and indeed the median width was 47 nm, represented a 9% decrease in median size (P = <0.05). The IQR was between 40 and 55 nm. This suggests that chromatin associated with different histone modifications represent different chromatin structures at the level of nucleosome clusters.

### The clusters are in mutually exclusive chromatin state domains

To more closely evaluate the domains of chromatin state along the Y loops, different histone modifications representative of active and inactive state, H3K36me3 and H3K27me3 respectively, were co-labelled, and imaged using STORM (Fig.8).

**Figure 8:**
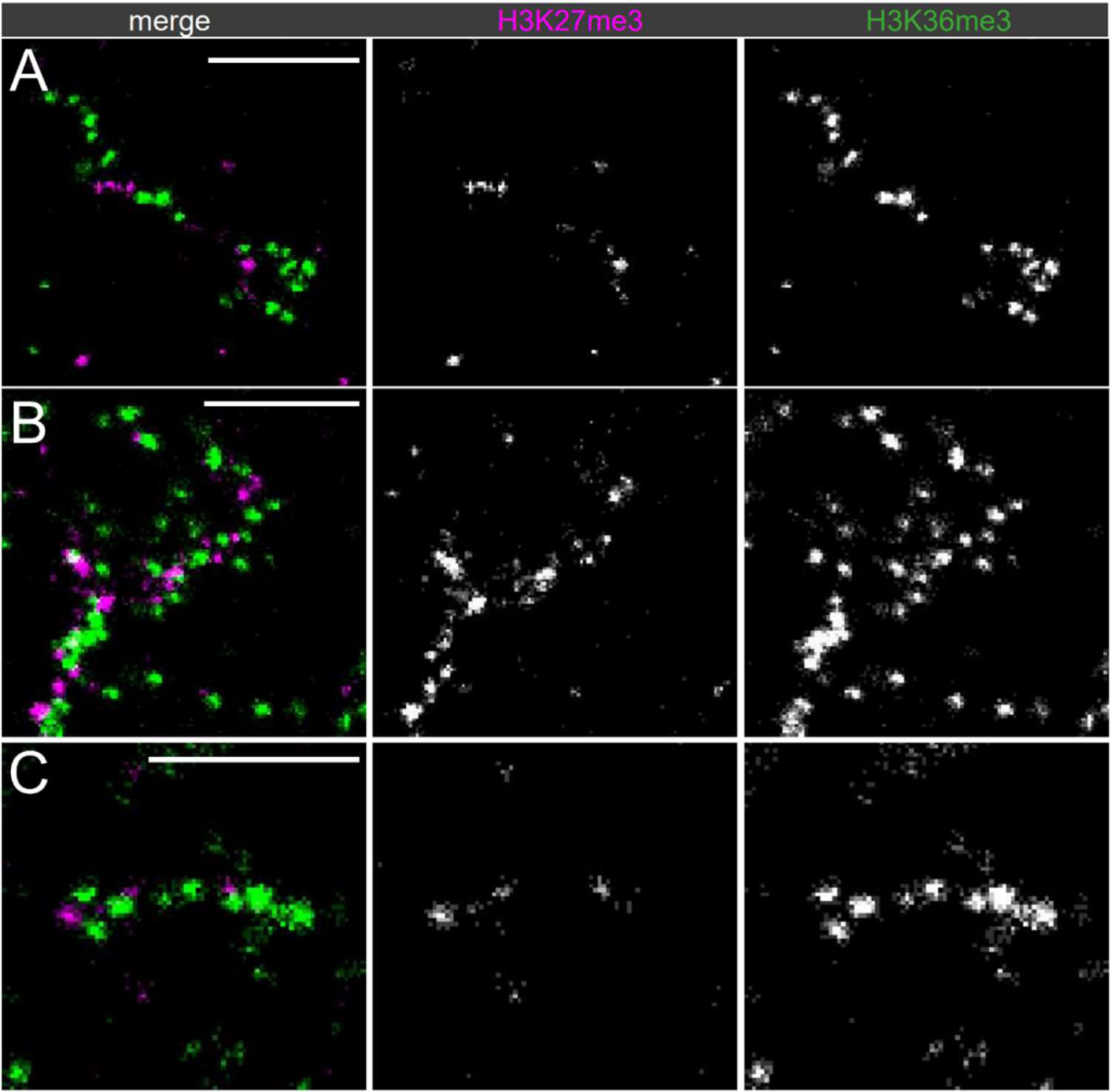
Dual-labelled super-resolution STORM images of the Y loops: H3K27me3 and H3K36me3. A-C) different examples are shown from three different nuclei labelled for H3K27me3 (magenta) and H3K36me3 (green). The scale bars are 1 µm.

Although both modifications can be seen along the same chromatin fibre nearby to one another, surprisingly, the chromatin clusters themselves appear to be mostly mutually exclusive for either modification. There is very little visual evidence of chromatin clusters that share two of the different types of modification together. First the apparent exclusivity hypothesis was tested and confirmed via the computation of the mark connection function, which demonstrated a lack of overlap of H3K27me3 and H3K36me3 localisations (Fig. 9C). This analysis also suggests that at distances smaller than 200 nm there is a distinct repulsion.

**Figure 9:**
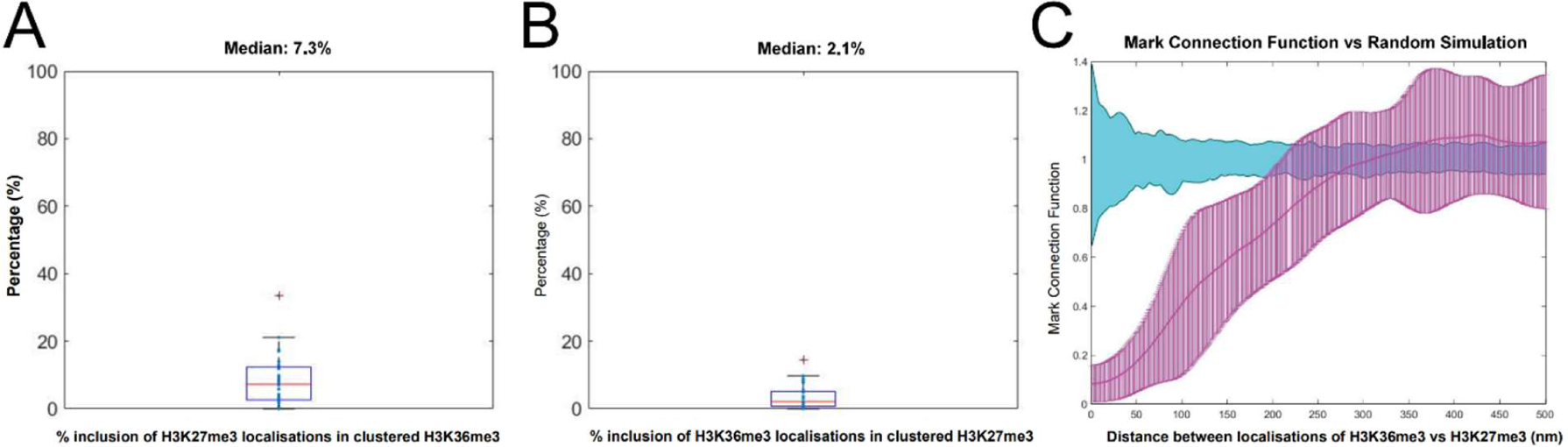
The exclusivity of H3K36me3 and H3K27me3 localisations. A) clustering the H3K36me3 localisations and quantifying the % of overlap of H3K37me3, and B) clustering the H3K37me3 localisations and quantifying the overlap of H3K36me3 localisations. C) mark connection function (magenta) vs randomly simulated data (cyan) was performed on H3K27me3 and H3K36me3 localisations along the Y loops to give confidence to the results with a more untargeted analysis. Values around zero show the chance of overlap between H3K27me3 and H3K36me3 localisations is random, those above zero show an attraction rule and those below zero show a repulsion rule. A strong repulsion between H3K27me3 and H3K36me3 is shown at distances below 200 nm.

To further quantify the exclusivity, both H3K27me3 and H3K36me3 were clustered, and the % of localisations within 50 nm radius of the cluster centres (as an approximation of average cluster width) was quantified (Fig 9). A median of 7% overlap existed between clusters of H3K36me3 and localisations of H3K27me3, demonstrating a clear exclusion of the two along a single Y loop chromatin fibre. Clustering H3K27me3, a median of 2% of H3K36me3 localisations were within 50 nm of the cluster centres.

This analysis makes some baseline assumptions, therefore in order to provide more confidence in this result, a general exclusivity quantification was undertaken using the mark connection function to further investigate this phenomenon, which also demonstrated a lack of overlap of H3K27me3 and H3K36me3 localisations (Fig. 9C). This analysis shows that at the cluster size level (< 200 nm) there is a distinct repulsion between H3K36me3 and H3K27me3, but above this distance is overlapping with random distribution, matching the data showing that clusters of both chromatin state can be next to one another, but not overlapping in one cluster. This has interesting implications for the mechanisms of how histone modifications spread in domains across chromatin, as this observation implies that individual chromatin clusters made up of many nucleosomes may form a “unit” of chromatin that can have different chromatin states, but that different chromatin states are not contained within the same nucleosome cluster.

There is evidence of regular switching between an active to an inactive chromatin state along one single chromatin fibre. In Fig. 8A, from left to right there is a continuous run of approximately 6 chromatin clusters labelled with H3K36me3, then a domain of a few clusters of H3K27me3, which switches back to a domain of H3K36me3 again. As previously described, this supports the findings that there are whole domains of different chromatin states along one fibre. However, at STORM resolution individual clusters along the Y loop are able to be visualised, and the extent of the proximity of these domains, and their apparent sharp boarders at chromatin cluster edges is striking.

This interspersion of different chromatin states can also be seen for H3K27me3 and H3K4me3 along Y loop chromatin (Fig. 10).

**Figure 10:**
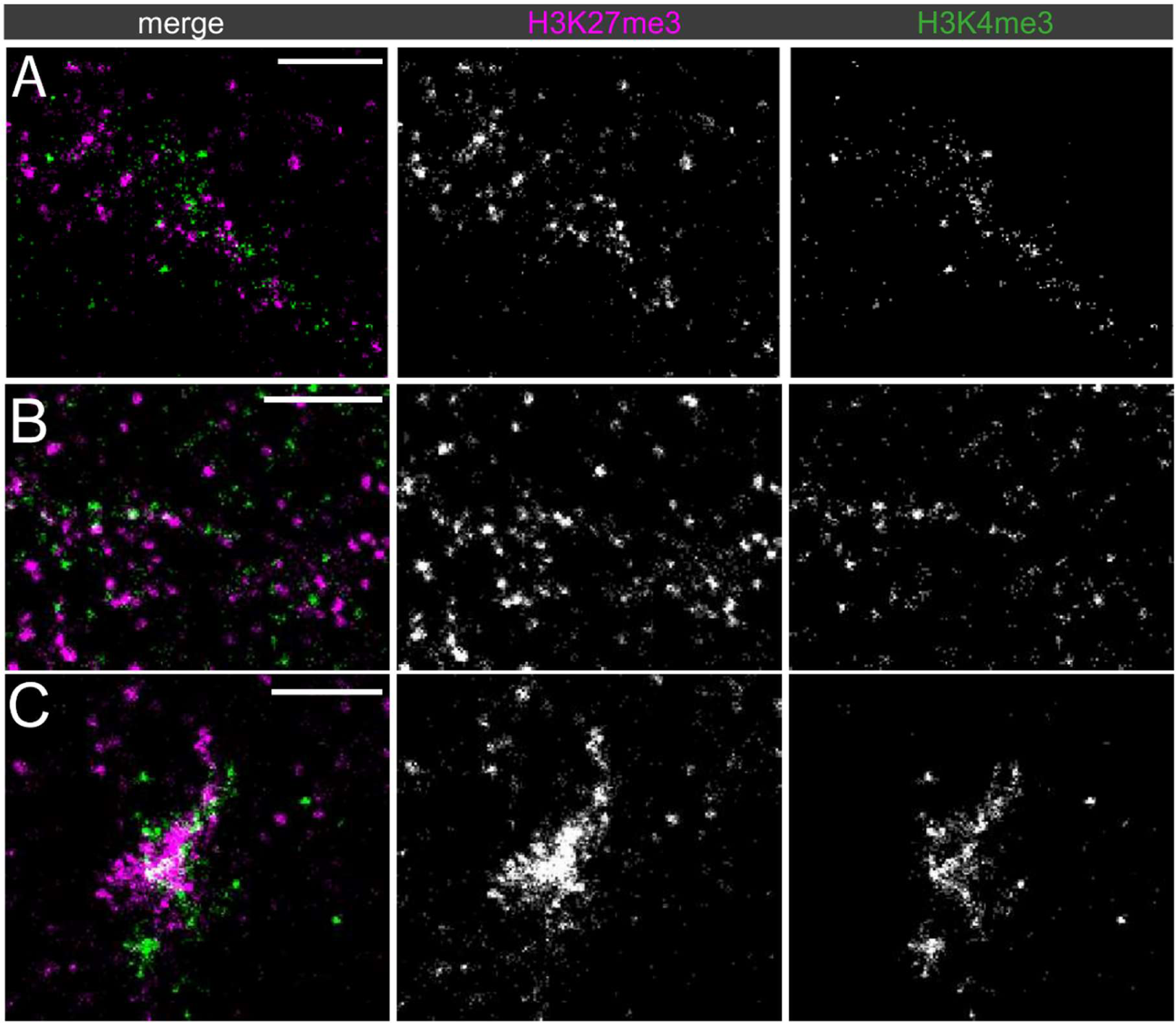
Dual-labelled super-resolution STORM images of the Y loops: H3K27me3 and H3K4me3. A-C) different examples are shown from three different nuclei labelled for H3K27me3 (magenta) and H3K4me3 (green). C) shows a dense region of chromatin on the periphery of the nucleolus thought to be near the origin point of the Y loops, and is strongly enriched for both H3K27me3 and H3K4me3. The scale bars are 1 µm.

### H3K36me3 is associated with elongating polymerase

To assess the functional association of chromatin state with active transcription, RNA Polymerase II with the phospho-Ser2 modification on the C-terminal domain repeats (RPol-Pser2) that is associated with active elongation (45,46) was immunolabelled alongside both H3K27me3 and H3K36me3 (Fig. 11).

**Figure 11:**
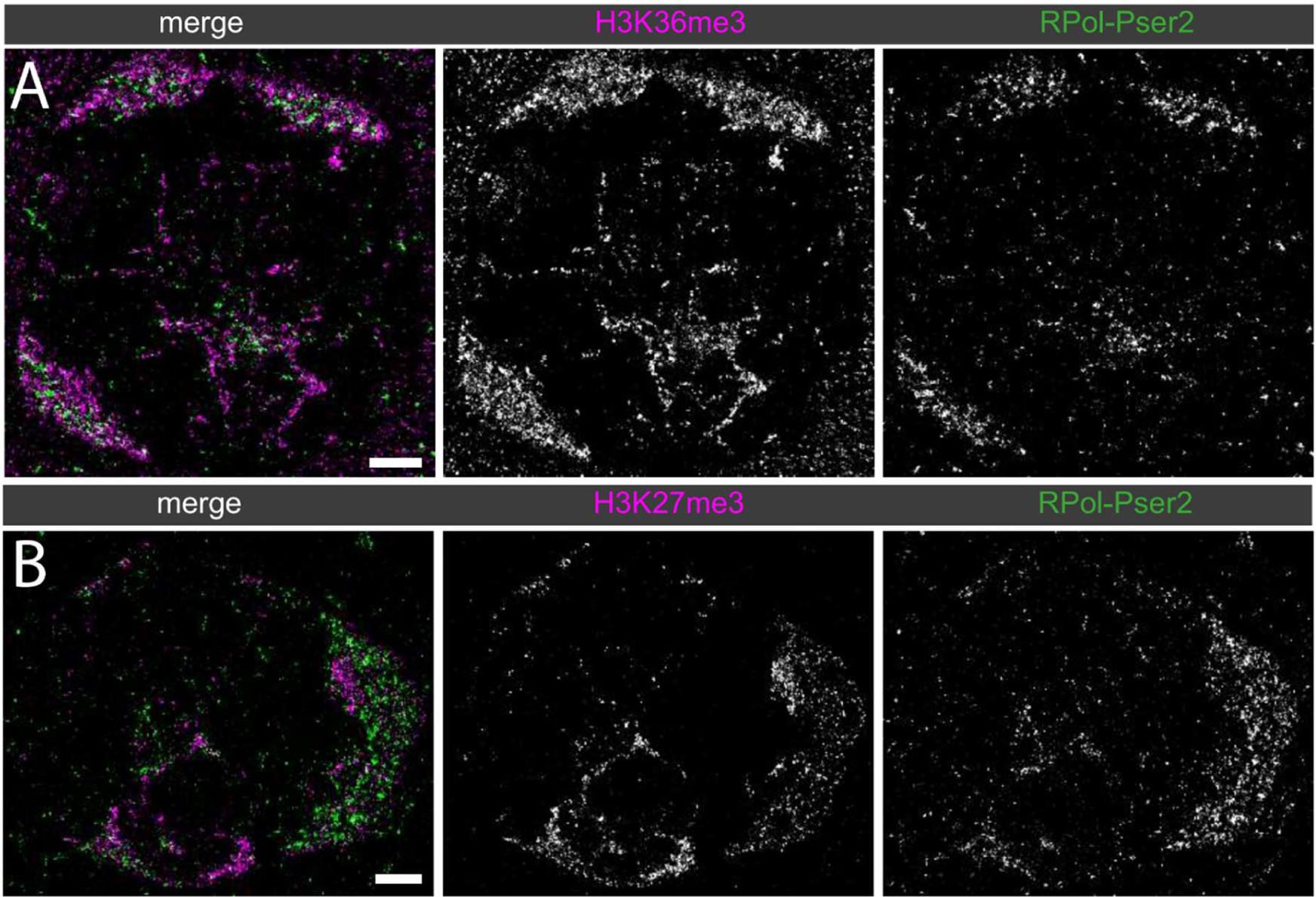
Dual-labelled super-resolution STORM images of primary spermatocyte nuclei labelled for histone marks with RPol-PSer2. A) H3K36me3 (magenta) and RPol-Pser2 (green) and B) H3K27me3 (magenta) and RPol-Pser2 (green). Scale bars are 2 µm.

RPol-Pser2 is still seen in association with chromatin labelled with H3K27me3, however, RPol-Pser2 appeared to be more closely associated with chromatin labelled with H3K36me3. To quantify this, the nuclear interior region of several cells labelled with H3K27me3 or H3K36me3 and RPol-Pser2 were cropped out. We computed the k-nearest neighbour distance for H3K27me3 and H3K36me3 localisations to RPol-Pser2 for k = 21, and the respective empirical cumulative distribution functions (Fig. 12A). Furthermore, we compared the median distance and the 90% quantiles for samples in the two cases (Fig. 12B). On average it was seen that H3K36me3 signal was significantly closer to RPol-Pser2 signal than to H3K27me3 along the Y loops (P = <0.05). A mark connection function confirmed this (Fig. 12C), indicating that RPol-Pser2 was significantly closer to H3K36me3 than H3K27me3 signal along the Y loops. A robustness test using a lower knn (5), and 50% quantiles confirmed the same trend of H3K36me3 having a closer association with RPol-Pser2 (see Supplementary Information).

**Figure 12:**
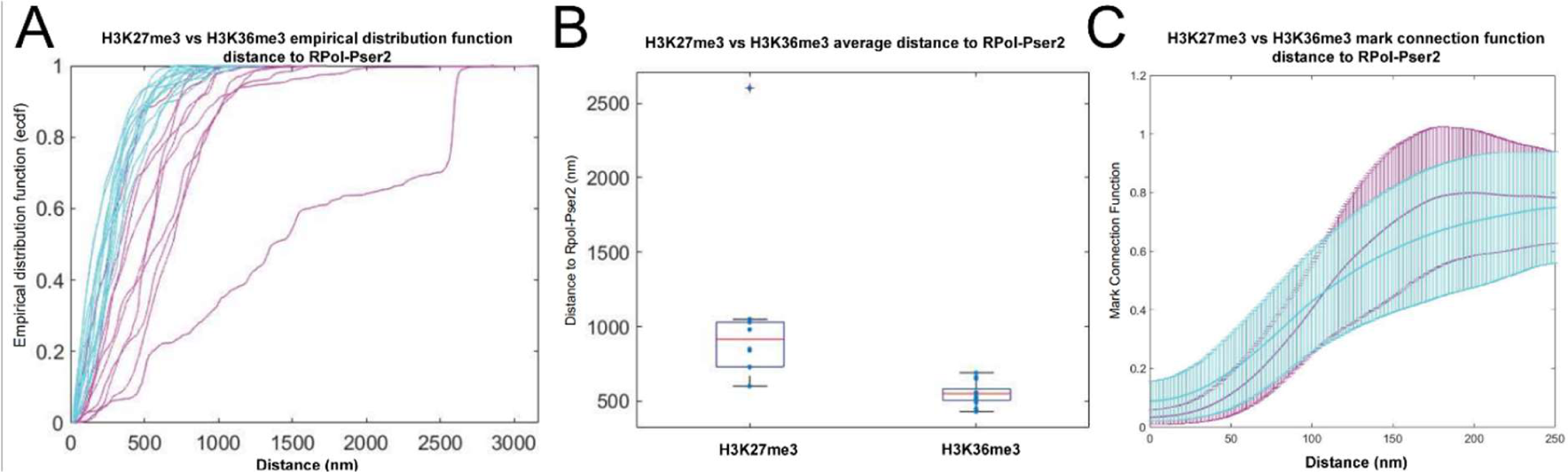
Quantification of the average overall distance between RPol-Pser2 and H3K36me3, or H3K27me3. A) individual ROIs containing the histone modifications and RPol-Pser2 along sections of the Y loop are plotted as individual lines using the empirical distribution function (ECDF); H3K36me3 in cyan, H3K27me3 in magenta. B) the ECDF values at 90% were pooled for both H3K27me3 and H3K36me3 and plotted on a box and whiskers plot showing that H3K36me3 was closer on average to RPol-Pser2 signal. C) the mark connection function was used as an alternative method in a more unbiased fashion, and also showed that RPol-Pser2 is closer to H3K36me3.

In specific examples of the association between H3K36me3 labelled Y loop chromatin and RPol-Pser2 it can often be seen that RPol-Pser2 is often associated on the edge of a domain of H3K36me3 (Fig. 13). This would align with a function of the enzyme SET2 that is responsible for catalysing the H3K36me3 modification, which may modify a region of chromatin as polymerase transcribes it, and the modification remains for a period of time behind the elongating polymerase. This is schematically represented in Fig. 13D.

**Figure 13:**
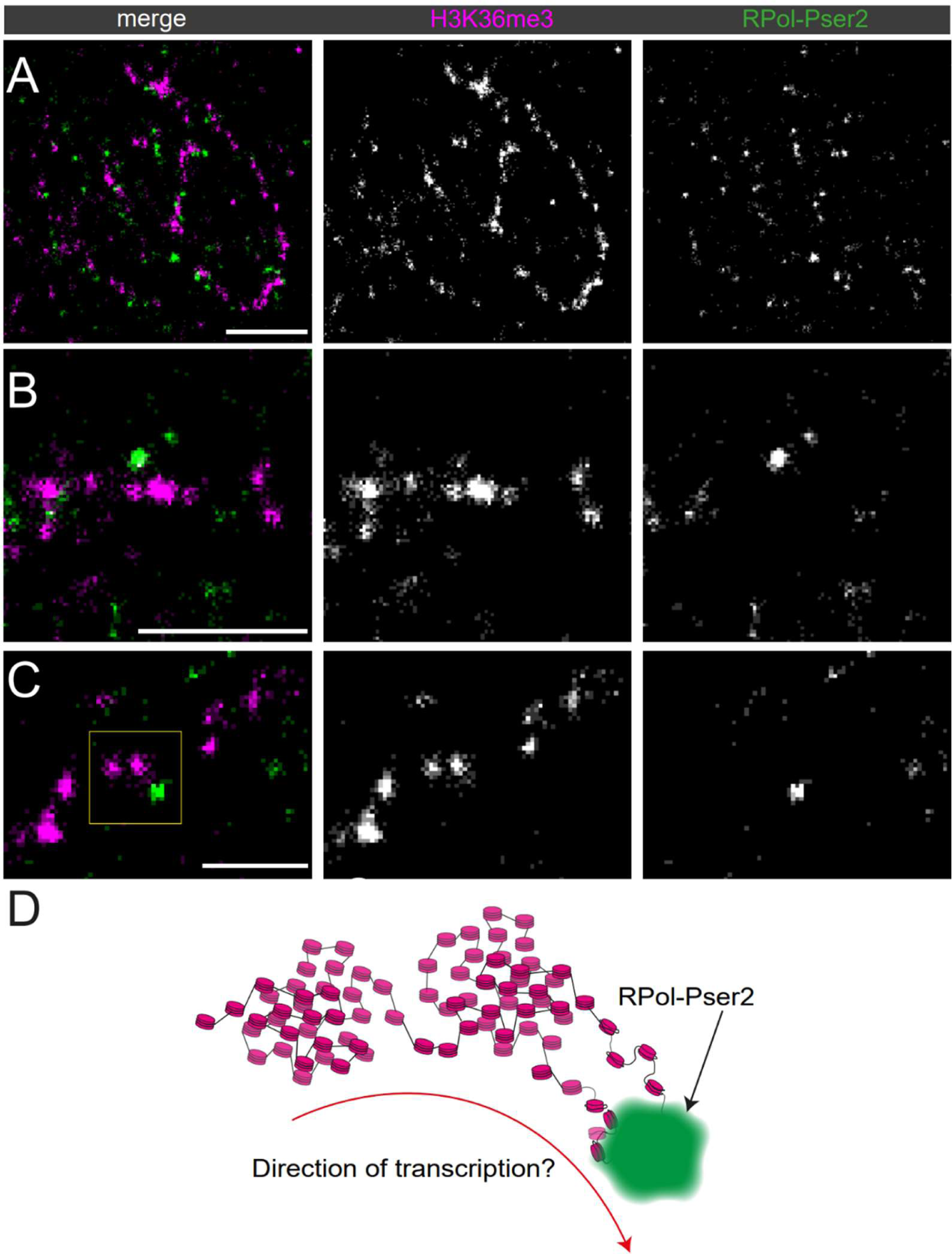
Dual-labelled super-resolution STORM images of the Y loops labelled for H3K36me3 (magenta) and RPol-Pser2 (green). A-C) different examples are shown from three different nuclei. D) schematic of selected area from C), indicated with a yellow box. The scale bars are 1 µm in A) and B), and 500 nm in C).

## Discussion

The model system of the Y loops in *Drosophila* spermatocytes has enabled us to examine the distribution of histone modifications along active transcription loops and to investigate the links between histone modifications and chromatin architecture. For the histone modifications examined, the ‘active’ marks of H3K4me3 and H3K36me3 and the ‘inactive’ mark H3K27me3, we find they are strikingly non-uniform in their distribution within transcription loops. The chromatin architecture of the Y loops is generally a chain of nucleosome clusters that extend out from the Y chromosome on the periphery of the nucleolus into the central nucleoplasm (33). The active histone marks are predominantly found on loop chromatin close to the nucleolus whereas H3K27me3 is present in domains more generally along the loops. Examining the link between histone marks and the structure of nucleosome clusters, we find that nucleosome clusters with different histone modifications clearly differ in their architecture. Remarkably, we also find a principle of exclusivity of histone marks at the level of nucleosome clusters. For example, although the H3K36me3 and H3K27me3 marks can be found on neighbouring clusters they show little, if any, overlap. This suggests that the nucleosome clusters are acting as a unit of chromatin state.

A transcription loop is a functionally dynamic structure with regions of transcription initiation, elongation, pausing, splicing and termination. These dynamics are apparently reflected in the distribution of chromatin marks and we observe independent chromatin stretches with either active or inactive marks as well as fibres where the active and inactive marks are more closely intermingled, indicating a briefer lifetime of the chromatin modifications (Figs. 4,8 and 10). Currently we can only speculate on the functional relevance of the observed distributions. The strong labelling of active marks on fibres close to chromosome mass of the Y chromosome around the nucleolus suggests a zone of enhanced transcriptional activity associated with the base of the loops and the presumed location of the loop promoters. The more sporadic distribution of active marks associated with Y loop fibres in the central nucleoplasm may also represent regions of transcription elongation and we find the H3K36me3 mark associated with active RNA polymerase (Fig. 13). The inactive mark H3K27me3 placed by the PRC2 complex is widespread over the active chromatin of the Y loops. In many regions it covers long domains but it is also present in regions of one or a few nucleosome clusters intermingled with active marks (Figs. 8 and 10) suggesting that it is highly dynamic. Although from genomic studies Polycomb complexes have been known to be associated with both silenced and expressed genes (47–49) we show here at the single cell, single gene level the association of the ‘inactive’ H3K27me3 mark with extended chromatin fibres undergoing transcription. This may reflect a balancing of inactive marks versus active transcription (reviewed in (50)), with PRC2 complexes targeting regions with less active transcription elongation. If so, it appears this can act at a very local level and potentially may provide a protective mechanism to control promiscuous transcription initiation from sites within chromatin in an extended “active” loop or to modulate the speed of transcription elongation.

Our studies indicate a clear link between specific histone modifications and the architecture of nucleosome clusters in the Y loops. We note that while Y loops form a diverse range of structures including condensed chromatin clumps, we generally focus on the extended fibres whose arrangement as chains of nucleosome clusters we analysed previously (33) and which are amenable to quantification. However, it is also worth noting that we did not observe an obvious connection between the inactive H3K27me3 mark and Y loop condensed chromatin clumps; some of these larger clumps were labelled for H3K27me3 but not all and the majority of the H3K27me3 was present on the extended fibres in chains of nucleosome clusters (Figs. 3 and 6). Also, even H3K4me3 was found occasionally in large dense chromatin accumulations (Fig. 1). We summarize our interpretations of the architectures of nucleosome clusters with different histone modifications in the schematic in Fig. 14. We interpret the sharper boundaries of the H3K27me3-marked clusters as representing more condensed clusters, and the fuzzier boundaries of the H3K36me3-marked clusters as representing a more open chromatin configuration with peripheral loops. We show in Fig. 13 that active RNA polymerase is associated with peripheral loops emanating from H3K36me3-marked clusters. The H3K4me3-marked chromatin appears to adopt a variety of architectures. From the STED images and analysis (Figs. 1 and 4) H3K4me3 labelling corresponded to regions of lower histone density, indicating relatively decondensed chromatin either between or peripheral to histone clusters. In the STORM images the H3K4me3 labelled chromatin formed small clusters and loops (Fig. 6) which, in double labelling with H3K27me3, appeared as small cluster or loop regions adjacent to H3K27me3-labelled clusters (Fig. 10). We have represented these various H3K4me3 potential architectures in Fig.14 as loops on the periphery of H3K27me3 clusters, as extended chromatin between nucleosome clusters, or as relatively decondensed small nucleosome clusters. The association of H3K4me3 with short regions of functionally accessible chromatin fits with the proposed role for H3K4me3 in transcriptional pause-release and elongation (40).

**Figure 14:**
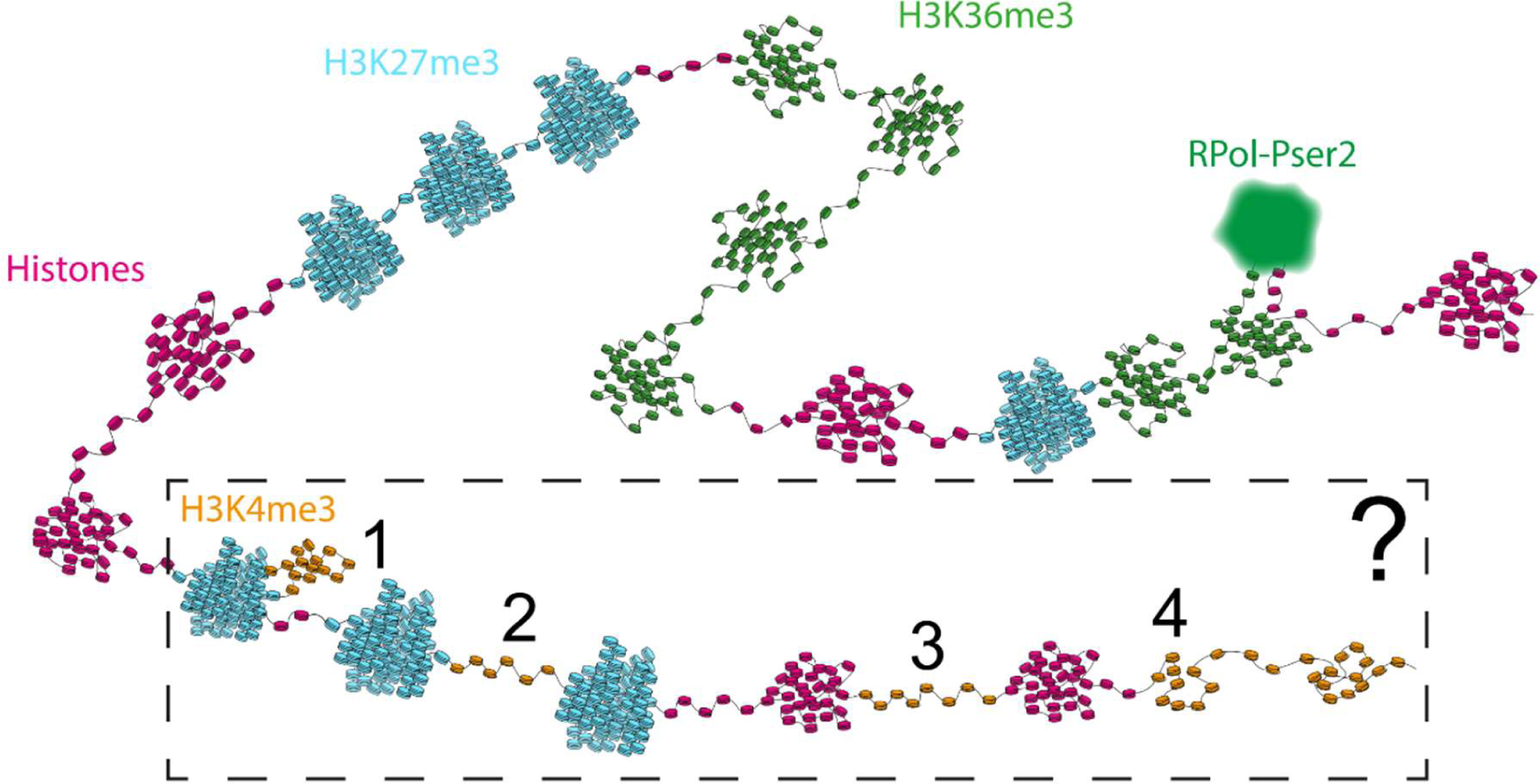
An overview schematic of the indicated architectures of the different modified chromatin along the Y loops. There are a few potential topological structures of H3K4me3, the options are shown within the boxed area. From left to right, H3K4me3 may appear as a looping region on the periphery of a H3K27me3 cluster (1), H3K4me3 may appear as a connecting region in between H3K27me3 clusters (2) or in between histone labelled clusters (3), or potentially as small clusters along a chromatin fibre (4). All of the structures indicated in this schematic are hypothetical based on the data presented in this work.

The link between histone modifications and specific nucleosome cluster morphology raises the question of whether the histone marks play a role in generating these different morphologies. We cannot answer this definitively however we note that each mark appears to be associated with a range of chromatin structures so, to this extent they do not determine specific structure. For example, although H3K36me3-marked clusters appear to have a specific morphology with peripheral loops of extended chromatin, the H3K36me3 mark is present both on the body of the cluster and on the peripheral loops. From this we conclude that the H3K36me3 mark does not specifically determine the extended chromatin structure of the peripheral loops. This raises a second question of whether histone marks play a role in generating the underlying general chromatin organisation of the Y loops as chains of nucleosome clusters. From the limited set of marks that we have studied, we see no evidence for this as both active and inactive marks are found associated with nucleosome clusters. This is also consistent with our previous results from transcription inhibition experiments showing that transcription is not required for chromatin clusters to persist (33). As chromatin can self-assemble into clusters in vitro in the absence of histone modifications (51), we favour a model where the underlying general chain of nucleosome clusters forms through a self-organising property of nucleosomes along a fibre. The histone modifications may then play a role in facilitating different cluster architectures in association with transcriptional processes.

A key observation from our studies is that we find very little overlap in the labelling of the different histone marks. We find this both between H3K4me3 and H3K27me3 (Figs. 5 and 10) and between H3K36me3 and H3K27me3 (Figs. 8 and 9). Detailed analysis of this for the case of H3K36me3 and H3K27me3 revealed that, although clusters labelled for either mark could be intermingled along fibres, individual clusters were exclusively labelled with one mark or the other. Thus, although the clusters contain several tens of nucleosomes there appears to be very little, if any, intermingling of H3K27me3 or H3K36me3-marked nucleosomes within clusters. We suggest this reveals an important principle that nucleosome clusters behave as units of chromatin state. Two processes may underlie this; firstly, the nucleosome clusters may facilitate the spread of histone marks through proximity and secondly, cross inhibition between histone marks may promote exclusivity. Proximity has been shown to be a key factor in the spread of histone marks, particularly H3K27me3 (52,53) and cross regulatory inhibition between H3K27me3 and H3K36me3 and H3K4me3 has been well documented (54,55). Individual nucleosome clusters of exclusive chromatin state may provide local hubs to facilitate transcription processes such as elongation and splicing.

## Materials and Methods

### Antibodies

Primary antibodies were: mouse anti-histone (core histones + H1, MabE71, Millipore), 1:1000; rabbit anti-RPol-PSer2 (ab238146, Abcam), 1:500; mouse anti-RPol-Pser2 (MA5-23510 ThermoFisher), 1:500; mouse H3K27me3 (ab6002, Abcam) 1:200; rabbit H3K27me3 07-449 (Millipore) 1:1000; rabbit H3K36me3 (ab9050) 1:500; rabbit H3K4me3 (ab8580). Secondary antibodies for STORM were from Invitrogen and used at 1:1000; goat anti-mouse Ig-Alexa Fluor 647 (A-21235), goat anti-rabbit Ig Alexa Fluor 488 (A-11008) and goat anti-rabbit Ig Alexa Fluor 568 (A-11011); for STED, Star580 (rabbit) (Abberior) 1:400; StarRed (mouse) (Abberior) 1:400.

### Spermatocyte Immunolabelling

Testes of 0–5 day old male w1118 *Drosophila melanogaster* were dissected in PBS. The primary spermatocytes were isolated via gentle pipetting following collagenase digestion (Sigma-Aldrich C8051, 5 mg/ml in PBS for 5min at room temperature) of the testes sheath, and fixed in 4% formaldehyde/PBS for 20min at 37°C. After washing with PT (PBS/0.01 % Tween 20) the primary spermatocytes were filtered (Partec 04-004-2327) and seeded onto 35 mm high μ-dishes (Ibidi) (STORM) or 13 mm No1.5h coverslips (STED and LSCM) for 30min at room temperature. The cells were blocked and permeabilized overnight (STORM) or for 2h (STED and LSCM) (1% Roche Western Blotting Reagent (WBR), Merck; 0.5% Triton X-100 in PBS). Following immunolabelling for 2h at room temperature with antibodies diluted in PBS/1 % WBR with 3x 20 min PT washes after each antibody incubation, the cells were fixed (4% formaldehyde) for 20min at room temperature and stored in PBS at 4°C.

### Microscopy

STED imaging was performed on an Abberior microscope with an Olympus IX83 frame using a 100x NA1.4 oil objective (UPLSAPO100XO) with excitation lasers 561 nm at 100 % and 640 nm at 9 % and a 775 nm STED laser at 8 % power with a dwell time of 10 µs, a pinhole of 1.00 AU and a pixel size of 20 nm and a Δz of 500 nm. Channels were acquired sequentially after each line scan. The spectral detection ranges were 590-630 nm and 650-765 nm for Abberior Star580 and Star Red, respectively. The STED laser was aligned using fluorescent beads. LSCM imaging was performed on a Leica SP8 with a 63x NA1.4 oil objective (HC PL APO CS2) with a GaAsP HyD detector and 561 nm and 633 nm laser lines in sequential mode, 16x line averaging, the pinhole set at 1 Airy Unit, with a voxel size of 80×80×300 nm. Detection bands were 555-625 nm and 645-760 nm. STED and LSCM data were recorded in 16 bit. For STORM cells were imaged using a Zeiss Elyra 7, at 30°C using a BP490-560/LP640 filter as described previously (33).

### Image analysis

Intensity peak distances along segmented line regions in STED data were calculated in Fiji using the FindPeaks plugin (56). Only fibre sections within single optical sections were chosen for analysis. Denoising for visual inspection only was performed with Noise2Void (57). STORM images were processed as described previously (33).

### Cross correlation analysis

For each ROI the position of maximum correlation lag was computed, the median displacement for H3K27 is close to zero (50 nm) meaning that the peaks and troughs of the two signals are well aligned, while for H3K4 the lag is 200 nm indicating that the location of high intensity values of the two signals tend not to match.

The lags for the two cases (H3K27 and H3K4) show significant difference, with the t-test giving a p-value of 0.0049. Non-significant correlation coefficient values were discarded (the cut-off values of +/− 1.96/√*n* were used for 5% significance).

### Exclusivity Analysis

In dual-labelled STORM images of H3K27me3 and H3K36me3, two parallel analyses were carried out to avoid bias due to the different abundance of the localisation from each histone modification. The localisations from one histone modification were clustered (MeanShift, sigma = 50 nm, minimum localisations in a cluster = 15), and the distances of localisation from the other modification to the nearest cluster centre were calculated. An average cluster width was assigned as 50 nm, therefore, if a localisation from the other class was within 50 nm of a cluster centre of the other modification class, this was considered an overlapping signal. In order to test the exclusivity hypothesis, mark connection function was also used (58). The mark connection function measures the probability that two points at distance r belong to different histone modification classes. A small (close to 0) mark connection function indicates that the two histone modification classes are more exclusive to each other than expected by chance, a high value indicates that the classes are more overlapping than expected by chance. This was compared to the random case – with expected mark connection function close to 1, simulated via random allocation of classes to points.

### Distance of H3K27me3 vs. H3K36me3 to RPol-Pser2 Analysis

Two datasets were considered for this analysis, dual-labelled H3K27me3 and RPol-Pser2, and dual-labelled H3K36me3 and Rpol-Pser2. The k-nearest neighbour distance of RPol-Pser2 localisations to the histone modification is less affected by false localisations. The parameter k was selected such that it is high enough to avoid noise but low enough so that the k’th neighbour is within the closest cluster (the accuracy being within the radius of the cluster). The knn distance from H3K37me3 and H3K36me3 localisations to RPol-Pser2 were therefore calculated for k = 5, 9, and 21 for robustness (see Supplementary Information) and the respective empirical cumulative distribution functions quantified. The median distance was calculated, and the 90% quantiles for samples in the two cases were compared using a t-test.

## Acknowledgements

We thank Martin Lenz (Cambridge Advanced Imaging Centre) for expert microscopy support.

## Author Contributions

RW conceived and supervised the project; MLB, VD and SAK carried out the experiments, MLB, SAK and LM carried out the analysis. MLB, SAK, LM and RW wrote the manuscript. All authors read and approved the final manuscript.

