## Supplemental Figures 1 & 2 for "Transcriptionally active chromatin loops contain both ‘active’ and ‘inactive’ histone modifications that exhibit exclusivity at the level of nucleosome clusters"

### Supplementary Information

#### 1. STED vs. Confocal

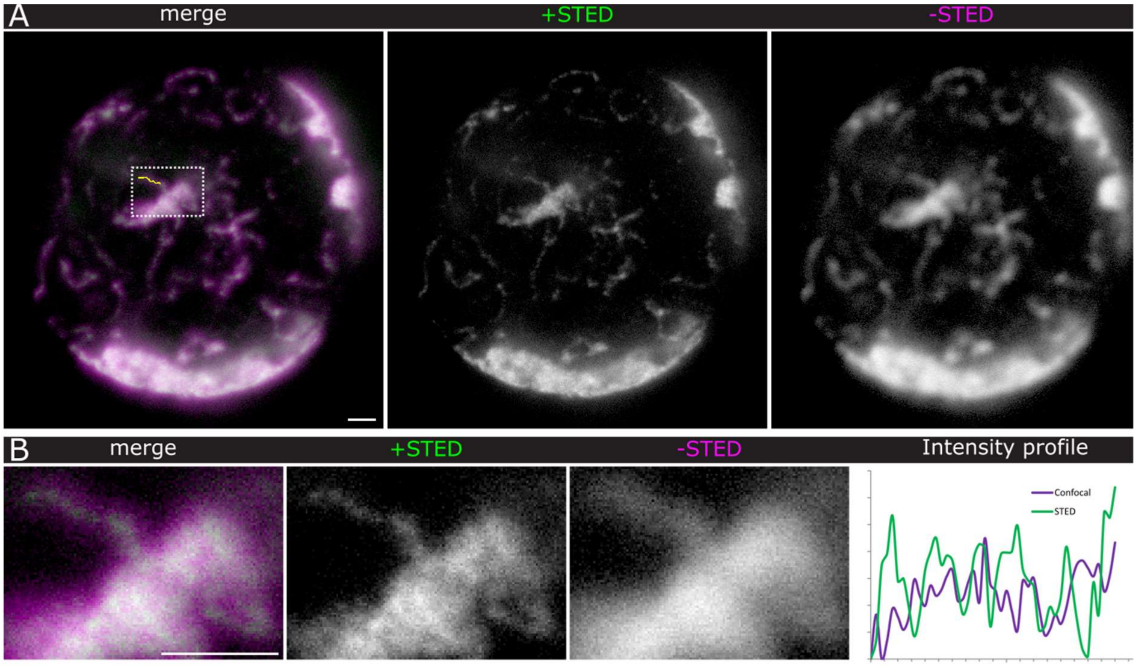


**Supplementary Figure 1**: Histones STAR-Red, single optical section, gamma=0.5, Scale bars = 1 µm; A) overview, B) detail boxed in A). Plot is the intensity profile (of the raw data (minus the minimum value)) along the line region indicated in A, merge. X-axis in µm, y-axis AU. Note the clearer peaks in the plot with STED imaging.

#### 2. Association of chromatin marks H3K36me3 and H3K27me3 with active RNA Polymerase

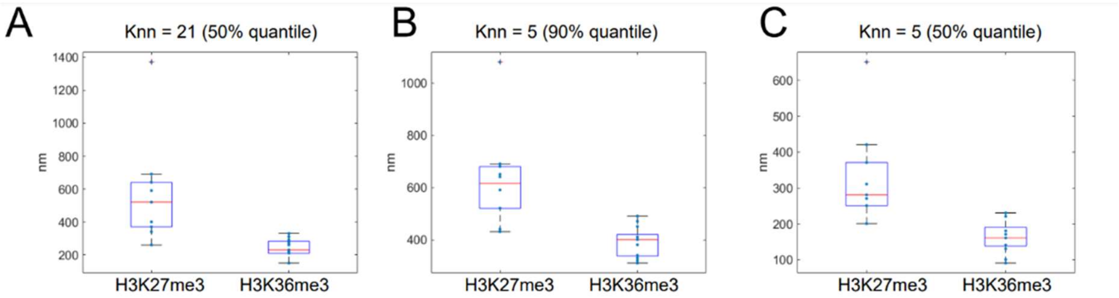


**Supplementary Figure 2**: A) To test whether the quantile selection was affecting the relationship between the histone modifications and RPol-Pser2, 50% was used, the closer association between H3K36me3 and RPol-Pser2 was still seen. B) To test whether the knn selection was affecting the relationship, a lower knn 5 was used with both B) 90% quantile, and C) 50% quantile. In all conditions, H3K36me3 had a lower distance to RPol-Pser2 localisations.
